# Cortico-brainstem mechanisms of biased perceptual decision-making in the context of pain

**DOI:** 10.1101/449652

**Authors:** K. Wiech, F. Eippert, J. Vandekerckhove, J. Zaman, K. Placek, F. Tuerlinckx, J.W.S. Vlaeyen, I. Tracey

## Abstract

Perceptual decision-making is commonly studied using stimuli with different physical properties but of comparable affective value. Here, we investigate neural processes underlying human perceptual decisions in the affectively rich domain of pain using a drift-diffusion model in combination with a probabilistic cueing paradigm. This allowed us to characterize a novel role for the dorsolateral prefrontal cortex (DLPFC), whose anticipatory responses reflecting a decision bias were dependent on the affective value of the stimulus. During intense noxious stimulation, these model-based anticipatory DLPFC responses were linked to an engagement of the periaqueductal gray (PAG), a midbrain region implicated in defensive responses including analgesia. Complementing these findings on biased decision-making, the model parameter reflecting sensory processing predicted subcortical responses (in amygdala and PAG) when expectations were violated. Our findings highlight the importance of taking a broader perspective on perceptual decisions and link decisions about pain with subcortical circuitry implicated in endogenous pain modulation.

## Introduction

Expectations critically shape the way we perceive pain: the same noxious input is, for instance, experienced as significantly more intense when we expect it to be of a high intensity (Atlas et al., 2012; Bingel et al., 2011; Keltner et al., 2006; Ploghaus et al., 2001; Yoshida et al., 2013). Such effects are often thought to be accompanied by changes in somatosensory processing. Using a model-based approach employing the drift diffusion model (Ratcliff, 1978; Vandekerckhove et al., 2011), we recently investigated whether the influence of expectations on pain perception could also be rooted in biased perceptual decision-making (Wiech et al., 2014). A direct comparison of both hypotheses confirmed the dominant influence of biased decision-making: cues signaling a higher probability of either low-intensity or high-intensity stimulation introduced a significant decision bias (as indexed by the model’s starting point parameter) towards the expected stimulation intensity rather than a change in sensory processing (as indexed by the model’s drift rate parameter).

Brain imaging studies have begun to unravel the neural underpinnings of such biased perceptual decision-making (Domenech and Dreher, 2010; Heekeren et al., 2008; Mulder et al., 2012; Philiastides et al., 2011), with evidence linking activity in the dorsolateral prefrontal cortex (DLPFC) to a shift in starting point in the assumed evidence accumulation process (Mulder et al., 2014). However, these findings originate from studies using affectively neutral stimuli (e.g., moving dots) and might not be applicable to the affect-rich domain of pain, giving rise to two possible scenarios. If activity in DLPFC indeed directly reflects the degree to which a perceptual decision is biased, cue-induced activity should be positively related to a shift in starting point towards the expected sensation, irrespective of the affective value, i.e. whether low or high pain is expected. On the other hand, if these responses are sensitive to the affective value of a choice, cue-induced activity could show a different, affect-dependent relation with a shift in starting point. Based on previous neuroimaging studies on pain–which have specifically implicated DLPFC regions in the expectation of reduced pain (Atlas and Wager, 2014) – one would expect a positive correlation with decision bias if low pain is expected and a negative correlation with decision bias when high pain is expected. This latter possibility would call into question the domain-generality of the above mentioned pattern of DLPFC responses, but would sit nicely with the known DLPFC involvement – together with the rostral anterior cingulate cortex (rACC) and the periaqueductal grey (PAG)– in descending pain control (Atlas and Wager, 2014; Seminowicz and Moayedi, 2017; Tracey, 2010; Wager and Atlas, 2015).

In addition to biased perceptual decision-making, we also focus on altered somatosensory processing (indexed by a change in the model’s drift rate parameter), which we had previously observed in the ‘worst-case’ scenario of a high-intensity painful stimulus being unexpectedly received (Wiech et al., 2014). In line with the assumption of altered somatosensory processing during expectancy-induced pain modulation, a change in drift rate should lead to activation changes in brain regions linked to somatosensory processing as well as the amygdala, which adjusts the sensitivity of sensory cortices depending on the perceived threat of the incoming sensory information (Fast and McGann, 2017; Hadj-Bouziane et al., 2012; Pourtois et al., 2013; Rotshtein et al., 2009).

Here we explore the neural processes underlying both biased decision-making and altered somatosensory processing in the context of pain using functional magnetic resonance imaging (fMRI) in a sample of healthy volunteers.

## Methods

The experimental paradigm and procedures as well as the analysis of the behavioural data including computational modeling have been described in detail in a previous publication (Wiech et al., 2014; for an overview see Fig. 1). The data presented here are based on these findings but focus on the fMRI data acquired in the same experiment. Below we therefore only give a summary of the behavioural part of the methods before we provide detailed information on the acquisition and analysis of fMRI data.

**Figure 1.**
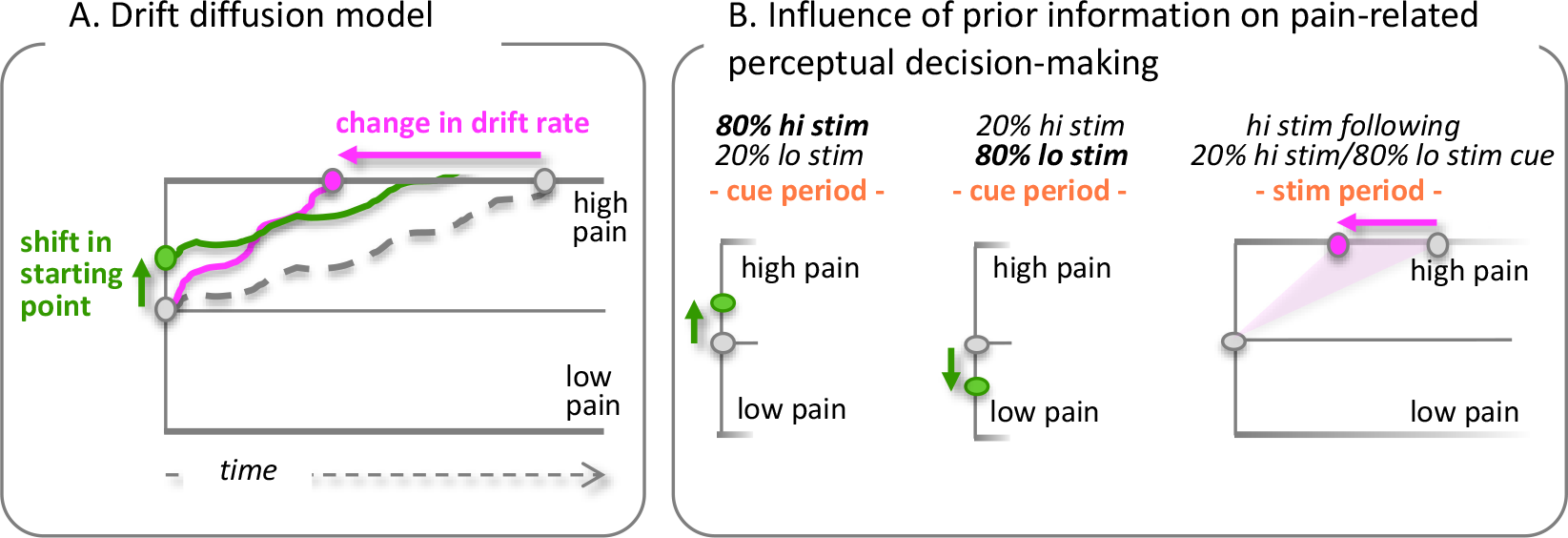
Overview of the Drift Diffusion Model (DDM) and previous behavioural findings. **(A)** The Drift Diffusion Model conceptualizes binary perceptual decisions as an inferential process in which sensory evidence is accumulated over time and the decision is made as soon as the upper or lower boundary is reached. The process is characterized by different parameters including the mean starting point (β) and the speed at which evidence is accumulated (i.e., the drift rate, δ). Prior information can bias the perceptual process through any of the parameters (unbiased process shown as dashed line). The graph separately depicts a shift in starting point (indicating a bias in decision-making; shown in green) and an increase in drift rate (indicating faster sensory processing; shown in magenta). (B) We previously demonstrated that a visual cue signaling an 80% probability to receive a high-intensity stimulation and 20% probability for low-intensity stimulation leads to a shift in starting point towards the high pain boundary relative to a 50/50 condition (left panel; Wiech et al., 2014). Similarly, a cue signaling a 20% probability for high-intensity and 80% for low intensity shifts the starting point towards low pain (mid panel). If a high-intensity stimulus is delivered following the presentation of cue signaling a 20% probability for high-intensity and 80% probability for low-intensity stimulation, an increase in drift rate is found (right panel). hi stim= high-intensity stimulation; lo stim= low-intensity stimulation; stim period= stimulation period.

### Participants

Twenty-two healthy volunteers (11 female; age *M*= 25.95 years, *SD*= 4.20) participated in the study. Sample size estimation was based on previous studies investigating expectancy effects on pain using fMRI (Atlas et al, 2010, N= 19) and studies exploring expectancy effects on perceptual decisions using a drift diffusion model in combination with fMRI (Mulder et al., 2012, N= 20; White et al., 2012, N= 24).

Participants were right-handed, fluent English-speaking, displayed normal pain thresholds at the site of stimulus application, and had normal or corrected-to-normal vision. According to self-report, no participant had a history of neurological or psychiatric disease or of chronic pain. Prior to involvement in the study, each participant gave full informed consent, and the study was approved by the local Research Ethics Committee (MSD-IDREC-C1-2013-106).

### Experimental paradigm

Using fMRI, we investigated brain responses during the anticipation and perception of high and low-intensity noxious electrical stimuli. The experiment was divided into four blocks, each consisting of 42 trials. On each trial, participants were presented with one of three visual cues: the white outline of a square, a triangle or a circle against a black background. One of these three cues indicated an 80% probability for high-intensity stimulation and a 20% probability for low intensity stimulation (‘80/20’ condition; note that the first number always refers to the probability of a high intensity stimulation, the second one to the probability of a low intensity stimulation). Another cue indicated a 20% probability for high-intensity stimulation and an 80% probability for low intensity stimulation (‘20/80’ condition). A third cue indicated a 50% probability for high-intensity stimulation and a 50% probability for low intensity stimulation (‘50/50’ condition). The three visual cues were randomly assigned to the 80/20, 20/80, and 50/50 conditions across participants, resulting in six different configurations of cue and condition pairings. In each configuration, the four blocks of trials were identical.

On each trial, either a high-intensity or a low-intensity stimulus was applied to the dorsal aspect of the left hand five seconds after presentation of the visual cue. Participants were instructed to indicate as quickly and accurately as possible whether they had received low-intensity or high-intensity stimulation by pressing one of two buttons with their right hand. Button-response contingencies were counter-balanced across participants. No feedback was provided regarding the correctness of the response on any trial. Decision accuracies and response times (RT; i.e. time between delivery of noxious stimulus and button pressing) were recorded. Each trial was completed by the presentation of a fixation cross for three, five, or seven seconds. The order of 80/20, 20/80, and 50/50 trials was pseudo-randomized with no more than two consecutive trials of the same type.

In preparation for the experiment, participants were first familiarized with the pairing between the three visual cues and the outcome probabilities and practiced providing their responses outside the MR scanner. Next, individual high and low-intensity stimulation levels were determined using a standardized calibration procedure. Subsequently, participants performed a discrimination test to ascertain whether individual levels of low-intensity and high-intensity stimulation were sufficiently different to allow for differential learning. If participants were able to categorize at least 80% of the trials correctly, they were positioned in the MR scanner and the experiment commenced.

### Electrocutaneous stimulation

Individual levels for high and low-intensity stimulation were determined for each participant using an ascending Method of Limits approach (Fruhstorfer et al., 1976). Intensities were rated on a Visual Analogue Scale (VAS) with the verbal anchor point “no pain” for the minimum intensity and “unbearable pain” for the maximum intensity. VAS ratings were transformed into a number between 0 and 10. Intensities rated as 2 were used for low-intensity pain, and intensities rated as 8 were used for high-intensity pain. Electrical stimuli were delivered using a commercial electric stimulation device (constant current stimulator DS7A; Digitimer, Hertfordshire, UK), delivering a 1ms monopolar square waveform pulse via a concentric silver chloride electrode attached to the back of the left hand. The average stimulation intensity across participants was 2.01 mA (*SD*= 1.54) for low-intensity stimuli and 8.20 mA (*SD*= 4.15) for high-intensity stimuli, which were perceived as significantly different (*t*(21)= 5.02; *p*< 0.001). The calibration procedure was first performed during preparation outside the scanner and both stimulation intensities were checked again and recalibrated if necessary once participants were positioned in the scanner and prior to each block to ensure constant pain levels throughout the experiment.

### fMRI data acquisition

Functional imaging was performed on a 3-Tesla MRI scanner (Siemens Verio, Siemens Medical Solutions) equipped with a 32-channel head coil. T2*-weighted echo-planar volumes with BOLD contrast were acquired at an angle of 30° to the anterior commissure-posterior commissure line to attenuate signal dropout in the orbitofrontal cortex (Deichmann et al., 2003) using a multiband EPI sequence (slice acceleration factor: 6; Moeller et al., 2010; Xu et al., 2013). A total of seventy-two transversal slices were acquired with an interleaved order for each volume, with an in-plane resolution of 2×2 mm and slice thickness of 2 mm (repetition time: 1300 ms; echo time: 40 ms; field of view: 212 x 212; flip angle: 66°). A whole-brain high-resolution T1-weighted structural scan (voxel size: 1×1×1 mm) was also obtained for each participant.

### fMRI data analysis: general aspects

Image processing and statistical analyses were performed using SPM12 (Wellcome Department of Imaging Neuroscience, London, UK; available at http://www.fil.ion.ucl.ac.uk/spm). Image processing consisted of slice timing (correction for differences in slice acquisition time), realignment (rigid body motion correction) and unwarping (accounting for susceptibility by movement interactions), coregistration (between EPI images and the skull-stripped T1 image), spatial normalization using the DARTEL toolbox, smoothing using an 8mm (full width at half maximum) isotropic three-dimensional Gaussian kernel and denoising using ICA-AROMA (Pruim et al., 2015).

For each participant, we constructed a design matrix that included three regressors for the anticipation phase (presentation of (i) the 50/50 cue, (ii) the 80/20 cue, and (iii) the 20/80 cue) and six regressors for the stimulus delivery phase (delivery of the low-intensity stimulation following (i) the 50/50 cue, (ii) the 80/20 cue, and (iii) the 20/80 cue and delivery of the high-intensity stimulation following (iv) the 50/50 cue, (v) the 80/20 cue, and (vi) the 20/80 cue), resulting in a total of nine regressors. All regressors consisted of delta functions convolved with the canonical hemodynamic response function. Six motion parameters derived from the realignment procedure were also included as regressors-of-no-interest. Model estimation was carried out using a robust weighted least squares approach (Diedrichsen and Shadmehr, 2005) and also included high-pass filtering (cutoff period: 128 s) and correction for temporal autocorrelations (based on a first-order autoregressive model).

Based on the first-level model, we defined contrasts of parameter estimates for each participant (see below) and subsequently explored the relation between individual modeling parameters (as derived using a Hierarchical Drift Diffusion model, HDDM; Vandekerckhove et al., 2011) and individual brain responses by means of regression analyses at the group level (for main effects during anticipation and stimulation period see Supplementary Tables 1 and 2). Modeling parameters considered here include the shift in starting point (indicating biased perceptual decision-making) and the change in drift rate (indicating altered sensory processing), always compared to the neutral 50/50 condition. Both parameters had been estimated for each participant as described in our previous publication (Wiech et al., 2014). In brief, we fitted the HDDM with four parameters (starting point, β; drift rate, δ; boundary separation, α; and non-decision time τ) using correct and incorrect trials. Drift rate and non-decision time parameters were allowed to differ between the six outcome conditions of the experiment (i.e., low-intensity stimulation following the ‘80/20’ cue, the ‘20/80’ cue or the ‘50/50’ cue and high-intensity stimulation following the ‘80/20’ cue, the ‘20/80’ cue or the ‘50/50’ cue). Boundary separation and starting point were only allowed to differ as a function of task instruction (‘80/20’ during anticipation, ‘20/80’ during anticipation or ‘50/50’ during anticipation) because both parameters are determined prior to stimulus delivery.

The regression analyses presented below are based on the significant behavioral effects reported in our previous publication (Wiech et al., 2014), which included a shift in starting point towards high pain when high pain is expected, a shift in starting point towards low pain when low pain is expected and an increase in drift rate when the high-intensity stimulation was unexpectedly applied (boundary separation and non-decision time were not investigated here, due to non-significant effects in our previous study). In all analyses, HDDM model parameters for the relevant condition were compared to those of the ‘50/50’ condition in which both outcomes were equally likely. These subject-specific differential model parameters were entered into simple regression analyses at the group level. In keeping with the differential HDDM modeling parameters, differential contrast images comparing the condition of interest with the non-informative ‘50/50’ condition were used as specified below. These analyses (as well as follow-up analyses on significant findings) are explained in detail below.

In all of our fMRI analyses, statistical inference was based on voxel-wise non-parametric permutation testing (Winkler et al., 2016) as implemented in SnPM (Statistical nonparametric Mapping; https://www2.warwick.ac.uk/snpm) using a threshold of p< 0.05 with family-wise error correction for both hypothesis-free whole-brain analyses and hypothesis-driven region-of-interest (ROI) analyses. ROIs for the current study included brain areas that are often ascribed a ‘pain modulatory’ role (DLPFC, rACC, amygdala, and PAG), as well as somatosensory brain areas that typically respond to noxious stimulation (thalamus, primary and secondary somatosensory cortex (SI, SII), posterior insula). Masks of these ROIs were derived from probabilistic atlases included with FSL (FMRIB’s Software Library; https://fsl.fmrib.ox.ac.uk/fsl/fslwiki/Atlases), all thresholded at 25%. In the case of cortical ROIs these masks were intersected with the participants’ average gray matter mask in order to exclude white matter regions of no interest. SI, SII, and posterior insula were based on the Juelich Histological Atlas (Eickhoff et al., 2005; SI (mask is a union of areas 1, 2, 3a, 3b): Geyer et al., 1999, 2000; Grefkes et al., 2001; SII (mask is a union of areas OP1, OP2, OP3, OP4): Eickhoff, 2005a, 2005b; posterior insula (mask is a union of areas lg1, lg2, ld1): Kurth et al., 2010), the thalamus mask was based on the Thalamic Connectivity Atlas (Behrens et al., 2003; mask is a union of all areas), the amygdala mask was based on the Harvard-Oxford Atlas (Desikan et al., 2006), the PAG mask was based on the PAG Atlas (Ezra et al., 2015; mask is a union of all areas), the DLPFC was based on the Dorsal Frontal Connectivity Atlas (Sallet et al., 2013; mask is a union of areas 9/46d and 9/46v) and the rACC was based on the Cingulate Orbitofrontal Connectivity Atlas (Neubert et al., 2015; mask is a union of areas 32d and 32pl). Note that we also provide unthresholded whole-brain maps of uncorrected p-values (again based on non-parametric permutation testing) for each tested effect at https://neurovault.org/collections/SHQGGEGD in order to aid interactive exploration of our data and to facilitate meta-analytic efforts.

### fMRI data analysis: anticipatory activity related to shift in starting-point

In order to test whether the DLPFC would show activation related to the degree to which the starting point had been shifted towards the high pain boundary, we performed a regression analysis on the differential imaging contrast between the presentation of the high pain cue and the non-informative cue (‘80/20 cue’ minus ’50/50 cue’) and used the relative shift in starting point between both conditions (‘80/20’ minus ‘50/50’) as the regressor-of-interest.

In order to test whether activation in the DLPFC also scaled with a shift in starting point towards low-intensity pain, we regressed the modeling parameters for the condition in which participants were cued towards low-intensity stimulation (‘20/80’ condition relative to the non-informative ‘50/50’ condition) against the equivalent imaging contrast (i.e., ‘20/80 cue’ minus ’50/50 cue’). Note that as starting point scores can vary between 0 (strongest possible bias towards low pain) and 1 (strongest possible bias towards high pain), the difference score between (numerically smaller) starting point indicating a bias towards low pain and a (numerically higher) starting point indicating no bias (as in the ‘50/50’ condition) would be negative. We therefore use the absolute value of the difference in starting point between both conditions (i.e., |‘20/80 cue’ minus ’50/50 cue’|). Although our hypotheses meant we focused analyses on the DLPFC (where we applied small-volume corrections separately for the right and left hemisphere), whole-brain analyses were additionally carried out, but revealed no significant results at a level of p< 0.05 FWE.

### fMRI data analysis: effects of anticipatory DLPFC activity

As explained in more detail in the results section, the previous analyses revealed a *negative* correlation between activation in DLPFC during cue presentation and the degree to which participants had shifted the starting point towards high pain. In order to investigate how this DLPFC engagement during the *cue period* translated into the *stimulation period* (i.e. when noxious stimuli were delivered), we performed two follow-up analyses. Individual parameter estimates were extracted from the identified DLPFC peak using the same contrast (i.e., ‘80/20 cue’ minus ‘50/50 cue’). These parameter estimates were entered as a regressor-of-interest in a group level regression analysis using the differential imaging contrast (‘80/20’ minus ‘50/50’) modeling activation during *stimulus delivery* (i.e. after the anticipation phase). Because the high pain cue could either be followed by low or high intensity stimulation, both trial types were analyzed separately: one analysis focused on trials in which high-intensity stimulation was delivered following the 80/20 cue (i.e., ‘80/20 cue followed by high-intensity stimulation’ minus ‘50/50 cue followed by high-intensity stimulation’) and another analysis focused on trials in which low-intensity stimulation was delivered (i.e., ‘80/20 cue followed by low-intensity stimulation’ minus ‘50/50 cue followed by low-intensity stimulation’).

The previous finding of a *positive* correlation between DLPFC activation and a shift in starting point towards low pain when low-intensity stimulation was the more likely outcome (‘50/50 cue’ minus ‘20/80 cue’; see Results for details) was followed up in a similar way: individual parameter estimates were extracted from the DLPFC peak identified in this analysis and entered as a regressor-of-interest into the imaging analysis testing for activity either during the unexpected delivery of high-intensity stimulation (‘20/80 cue followed by high-intensity stimulation’ minus ‘50/50 cue followed by high-intensity stimulation’) or during the expected delivery of low-intensity stimulation (‘20/80 cue followed by low-intensity stimulation’ minus ‘50/50 cue followed by low-intensity stimulation’).

Based on the tight link between DLPFC, rACC and PAG in descending pain modulation (Bingel et al., 2006; Eippert et al., 2009; Stein et al., 2012; Wager et al., 2004), we applied small-volume correction to the rACC and PAG in these analyses (whole-brain analyses were also carried out but revealed no significant results at a level of p < 0.05 FWE).

### fMRI data analysis: stimulation period activity related to an increase in drift rate

As reported in our previous publication (Wiech et al., 2014), the unexpected application of a high-intensity electrical stimulus led to an increase in drift rate. To test our hypothesis of whether brain regions involved in pain-related sensory processing and the amygdala reflected this change in somatosensory processing, we performed a regression analysis using the difference between the drift rate of the condition in which participants had been cued towards low pain, but received a high-intensity stimulus (‘20/80’ condition with high-intensity stimulation) and the neutral ‘50/50’ condition with high-intensity stimulation as a regressor-of-interest for the differential contrast image of both conditions (i.e., ‘20/80’ condition with high-intensity stimulation minus ‘50/50’ condition with high-intensity stimulation). Given our hypotheses, these analyses focused on somatosensory regions (thalamus, SI, SII and posterior insula) as well as the amygdala, where we applied small-volume corrections (whole-brain analyses were also carried out and revealed one significant response in the left hippocampus: x,y,z= −30,-14,-10; t= 6.49; p= 0.035 FWE).

### fMRI data analysis: connectivity changes related to an increase in drift rate

In order to investigate the functional connectivity of the drift rate related amygdala activation identified in the previous analysis we conducted a psychophysiological interaction analysis (PPI; Friston et al., 1997) using the left and right amygdala peaks as seed regions in two separate analyses. For each individual we first extracted the BOLD time-series from the peak voxel of the left and right amygdala activation identified in (x,y,z= −26,-8,-14 for left amygdala and x,y,z= 26,-2,-12 for right amygdala). Next, a PPI regressor was computed as the element-by-element product of the mean-corrected amygdala activity and a vector coding for the condition in which a high-intensity stimulus was unexpectedly applied compared to high-intensity stimulation following the non-informative cue (i.e. ‘20/80’ high pain minus ‘50/50’ high pain). The individual contrast images reflecting the interaction between this psychological variable (i.e., the difference between ‘20/80’ high pain and ‘50/50’ high pain) and the activation time course in the amygdala were subsequently entered into a second-level regression analysis with the individual difference in drift rate between both conditions (i.e., ‘20/80’ high-intensity stimulation minus ‘50/50’ high-intensity stimulation) as the regressor-of-interest. While our initial hypothesis was that the unexpected delivery of a high-intensity stimulation should alter the information exchange between amygdala and brain regions related to somatosensory processing (SI, SII, thalamus and posterior insula), we also explored the possibility that such a scenario could lead to enhanced connectivity between the amygdala and the PAG, as suggested by a large line of evidence from the fear conditioning literature (Johansen et al., 2010; McNally et al., 2011; Ozawa and Johansen, 2018; Tovote et al., 2016). We thus carried out small volume corrections for these regions, but also ran whole-brain analyses, which however revealed no significant results at a level of p< 0.05 FWE.

### Correlation between prior and drift rate

In order to explore the relationship between a bias in decision-making (reflected in the starting-point) and a bias in sensory processing (reflected in the drift rate), we calculated the Pearson correlation coefficient between both modelling parameters separately for each of the six conditions. Correlations with p< 0.05 (2-tailed) after correction for multiple comparisons were deemed significant.

## Results

Based on the computational modeling of decision accuracies and response times using a hierarchical drift diffusion model as described in our previous publication (Wiech et al., 2014), our interrogation of the fMRI data focused on three reported behavioural effects: the shift in starting point towards high pain during the expectation of high-intensity stimulation, the shift in starting point towards low pain during the expectation of low-intensity stimulation, the increase in drift rate when a low-intensity stimulation was expected but a high-intensity stimulus was unexpectedly delivered (Fig. 1), and follow up analyses based on results from these.

### Anticipatory activity related to shift in starting point

Analyses investigating the shift in starting point towards high pain did not provide evidence for a positive correlation, but instead revealed a negative correlation between the relative shift in starting point towards the high pain boundary and activation in the right DLPFC (x,y,z= 22,36,52; t= 4.35; p= 0.023; Fig. 2A,B). In other words, the more participants activated the DLPFC during the cue period when they were expecting the high-intensity stimulation (relative to the ‘50/50’ condition), the weaker was their decision-making bias towards high pain. When performing the same analysis for a shift in starting point towards low pain, we observed a positive correlation with right DLPFC activity (x,y,z= 48, 30, 34; t= 4.28; p= 0.025 and x,y,z= 46,22,42; t= 3.98; p= 0.043; Fig. 2C,D). So, the more participants activated the DLPFC during the cue period when they were expecting the low-intensity stimulation (relative to the ‘50/50’ condition), the stronger was their decision-making bias towards low pain. Together, these results stand in clear opposition to the pattern of DLPFC responses observed in affectively neutral decision-making scenarios, and instead suggest that they might be related to preparatory ‘protective’ function.

**Figure 2.**
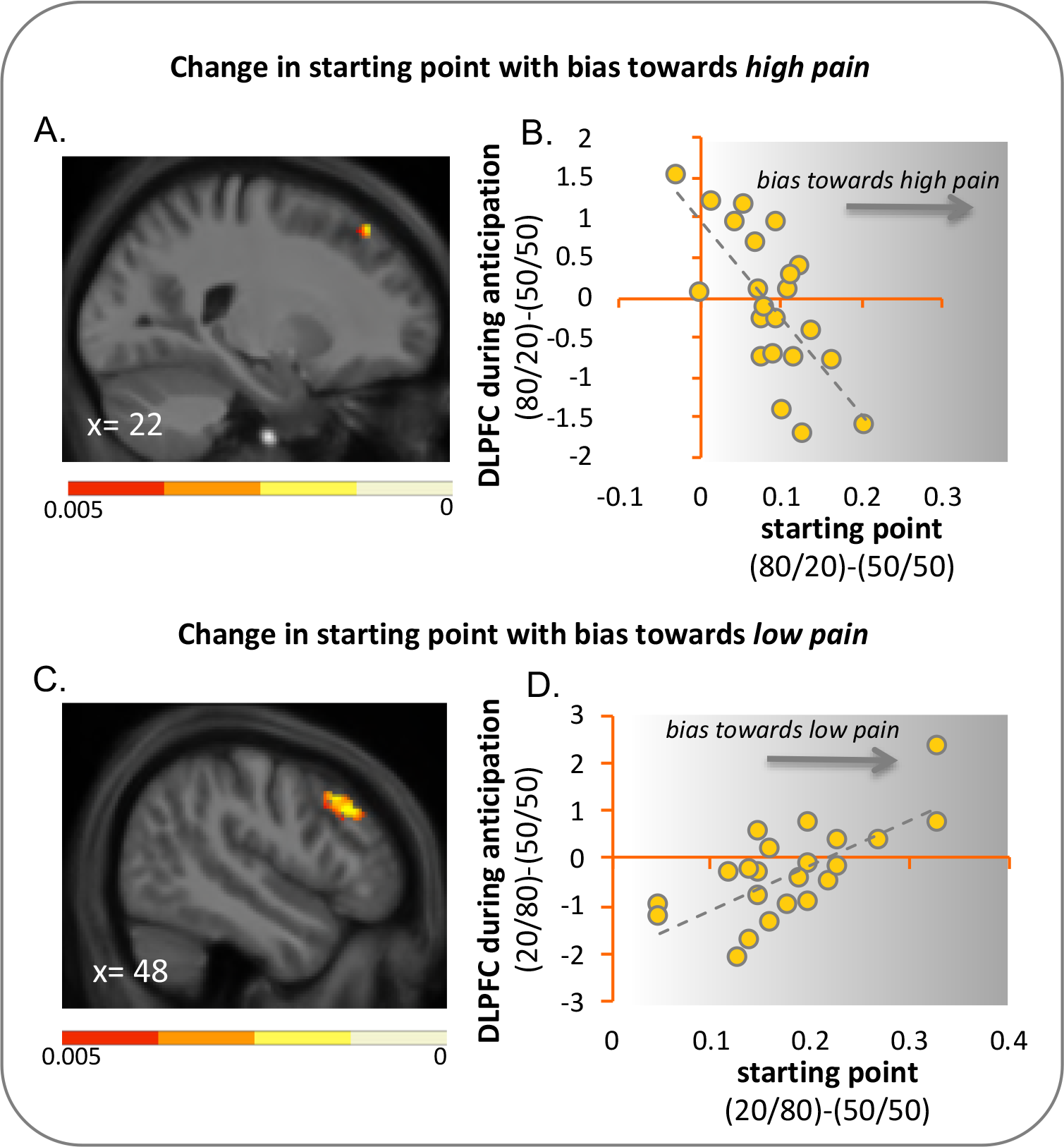
DLPFC activation related to a shift in starting point. **(A)** DLPFC activity exhibiting a *negative* correlation with change in starting point when high-intensity stimulation was the more likely outcome relative to the condition when both stimulation intensities were equally likely (‘80/20’ minus ‘50/50’; thresholded at p< 0.005 uncorrected for display purpose; overlaid on group mean T1 image masked by DLPFC region-of-interest). (B) Illustrative scatter plot showing the relationship between change in starting point (‘80/20’ minus ‘50/50’) and activation in the peak voxel of the DLPFC (‘80/20’ minus ‘50/50’). (C) DLPFC activation exhibiting a *positive* correlation with change in starting point towards the low pain boundary when low-intensity stimulation was the most likely outcome relative to the condition when both stimulation intensities were equally likely (‘20/80’ minus ‘50/50’; thresholded at p< 0.005 uncorrected for display purpose; overlaid on group mean T1 image masked by DLPFC region-of-interest). (D) Illustrative scatter plot showing the relationship between change in starting point (‘20/80’ minus ‘50/50’) and activation in the peak voxel of the DLPFC (‘20/80’ minus ‘50/50’). Note that the x-axis shows the absolute difference in starting point between both conditions (|’20/80’ minus ‘50/50’|).

### Effects of anticipatory DLPFC activity

In order to further explore the pain relevance of this affect-dependent anticipatory engagement of the DLPFC, we tested whether DLPFC activation during the *stimulus anticipation* (i.e. related to the shift in starting point) would predict activation in down-stream regions of the descending pain control system, namely rACC and PAG, during *stimulus application.* While we investigated both shifts in starting point (i.e. towards low pain and towards high pain) in combination with the actual delivery (i.e. delivery of high pain and delivery of low pain), we only observed a DLPFC-dependent recruitment of the descending pain control system during the expected delivery of high pain. In this case, the identified DLPFC activation during stimulus anticipation was positively related to PAG engagement during stimulus receipt (x,y,z= 2,-30,-10; t= 3.49; p= 0.016 [89% vIPAG, 32% lPAG] and x,y,z= −2,-28,-10; t= 3.48; p= 0.017 [58% vIPAG, 5% lPAG]; Fig. 3). Such a relationship (or its inverse) was neither observed when expectations where violated (i.e. delivery of low pain when cued about high or delivery of high pain when cued about low pain) nor when the expectation of low pain was confirmed. Interestingly, the observed PAG activation was localized to the ventrolateral PAG, i.e. the PAG column that is most strongly associated with descending analgesia.

**Figure 3.**
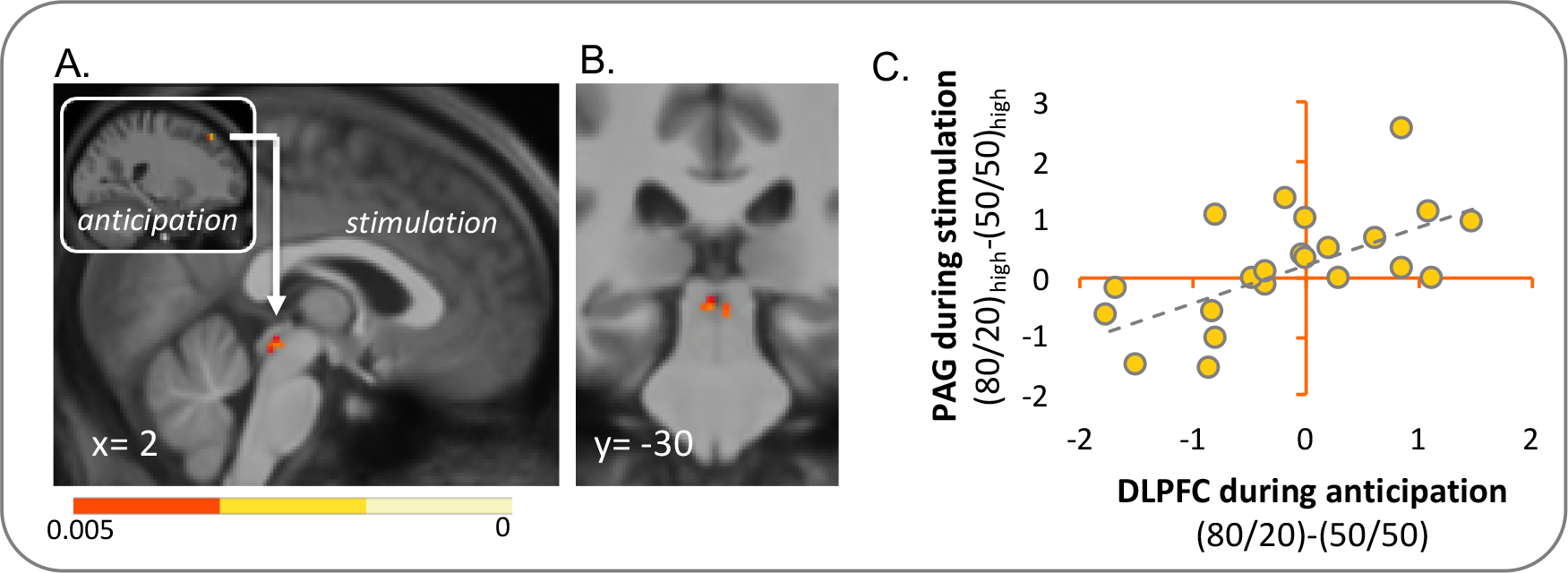
DLPFC-related PAG responses during the stimulation period. (A) Differential DLPFC activity (shown in inset) during the *anticipation* of high-intensity stimulation (relative to the ‘50/50’ condition) was used as a covariate in the analysis of activity during high-intensity *stimulation* following the ‘80/20’ cue (relative to the ‘50/50’ cue). DLPFC activity was extracted from the peak voxel identified in Analysis 1. (A) Sagittal view and (B) coronal view of the activation cluster in the PAG during delivery of the expected high-intensity stimuli (thresholded at p< 0.005 uncorrected for display purpose; overlaid on group mean T1 image masked by PAG region-of-¡nterest). (C) The illustrative scatter plot shows the positive correlation between differential (‘80/20’ minus ‘50/50’) anticipatory DLPFC activity and differential (‘80/20’ minus ‘50/50’) PAG activity during delivery of high-intensity stimulation across the sample.

### Stimulation period activity related to an increase in drift rate

Turning to changes in sensory processing, we found a significant association between an increase in drift rate and activation in the left and right amygdala (left: x,y,z= −26,-8,-14; t= 6.02; p= 0.002; right: x,y,z= 26,-2,-12; t= 4.16; p= 0.041; Fig. 4) when expectations were violated towards the worse. In other words, the stronger the amygdala responses, the higher the increase in drift rate during the application of high-intensity stimuli that followed a cue signaling delivery of a low intensity stimuli. Contrary to our hypothesis, such a relationship was only observed in the amygdala, but not in brain regions related to somatosensory processing (i.e. thalamus, SI, SII and posterior insula).

**Figure 4.**
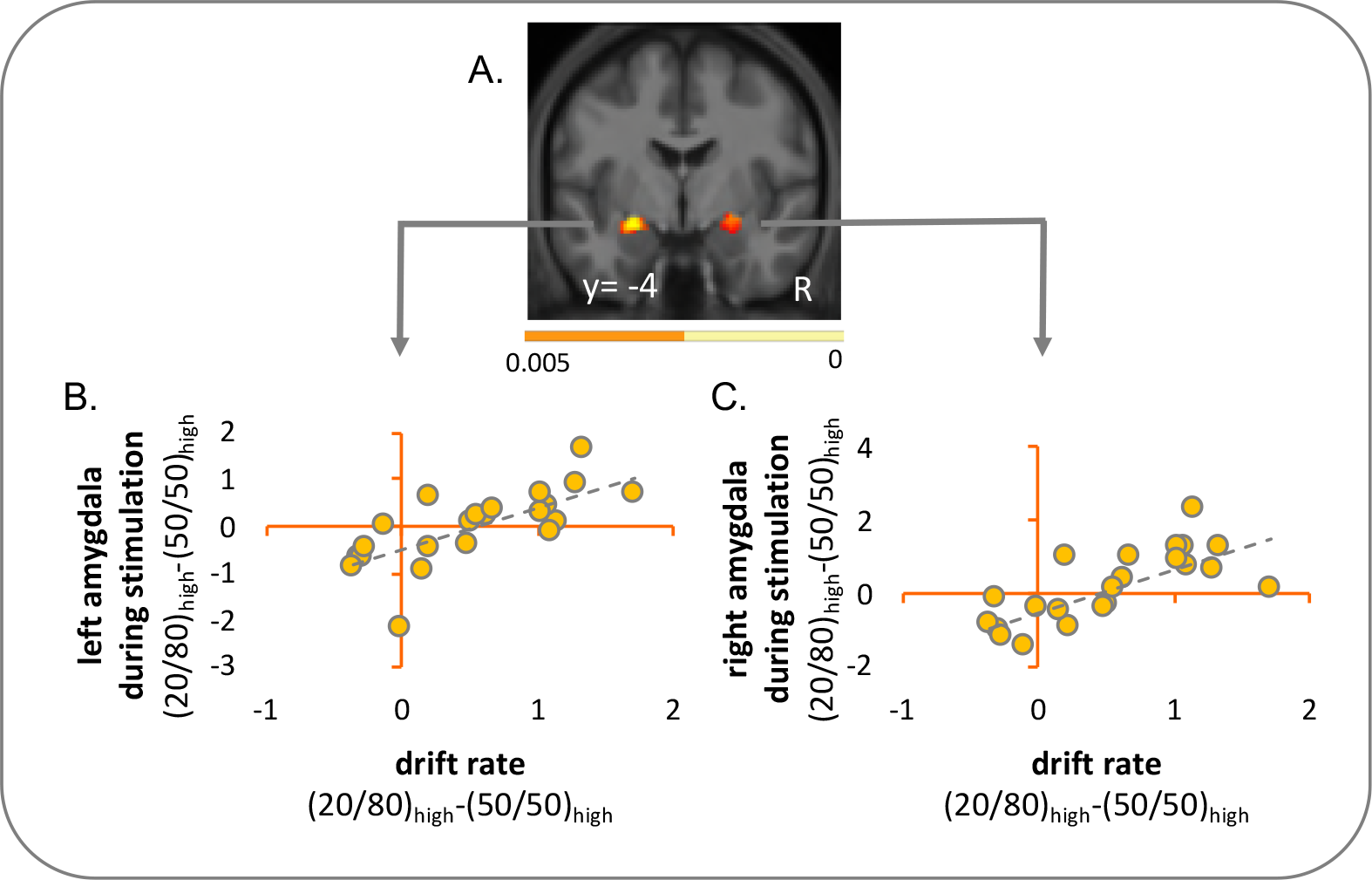
Drift-rate related activity in the amygdala during unexpected delivery of high-intensity stimuli. (A) Activity in left and right amygdala during the delivery of high-intensity stimulation following the presentation of the safe cue (‘20/80_high’) scaled with an increase in drift rate (both relative to the neutral ‘50/50’ condition; thresholded at p< 0.005 uncorrected for display purpose; overlaid on group mean T1 image masked by amygdala region-of-interest). Illustrative scatter plots from the amygdala peak voxels show the correlation between the condition difference in drift rate and activation in the left (B) and right (C) amygdala across the sample.

### Connectivity changes related to an increase in drift rate

In order to further explore the consequences of the drift rate related amygdala responses during the ‘worst case scenario’ (i.e. delivery of high-intensity stimulation when low-intensity stimulation was more likely), we carried out psychophysiological interaction analyses seeded in the amygdala. These analyses showed a context-dependent increase in functional connectivity between the left amygdala and the PAG (x,y,z= −2,-36,-10; t= 3.68; p= 0.010 [79% dmPAG, 11% dlPAG, 5%vlPAG, 5% lPAG]; x,y,z= 4,-28,-4; t= 3.51; p= 0.014 [68% lPAG, 21% vIPAG, 21% dmPAG, 16% dlPAG]; Fig. 5), but did not provide evidence for the hypothesized increase in connectivity with somatosensory brain regions (i.e., SI, SII, thalamus and posterior insula). Interestingly, the PAG responses we observed here exhibited a different spatial pattern compared to the ones reported above in relation with the DLPFC (mainly ventrolateral PAG), focusing more on dorsal and lateral aspects of the PAG.

**Figure 5.**
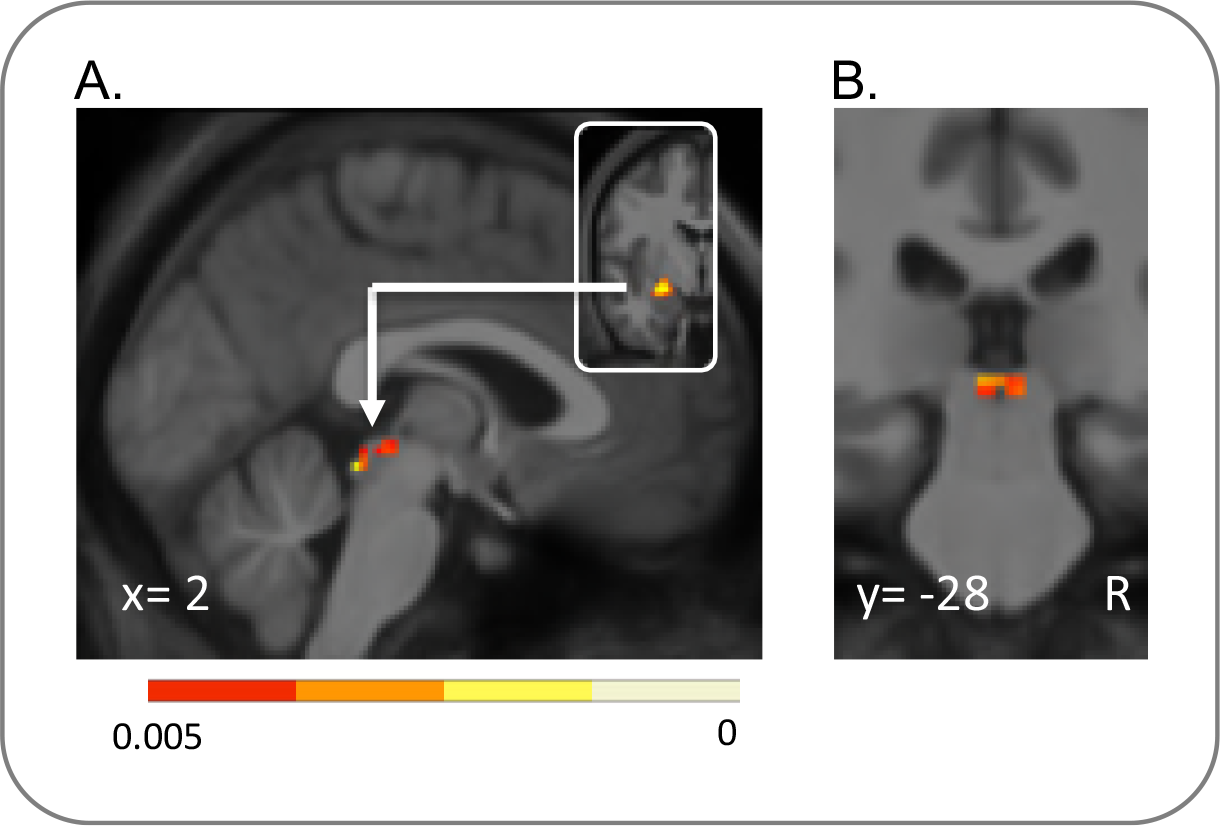
Drift-rate related functional connectivity between amygdala and PAG during unexpected delivery of high-intensity stimuli. (A) Sagittal and (B) coronal view of the PAG showing increased functional connectivity with the left amygdala (inset) during the unexpected delivery of high-intensity stimuli (relative to high intensity stimulation following the ‘50/50’ cue; ‘20/80_high’ minus ‘50/50_high’) depending on drift rate (‘20/80_high’ minus ‘50/50_high’). Thresholded at p< 0.005 uncorrected for display purpose; overlaid on group mean T1 image masked by PAG region-of-interest.

### Correlations between starting point and drift rate

Correlations between both model parameters were positive in all of the six comparisons but only reached significance in the condition when high pain was expected but the low intensity stimulation was delivered (‘20/80’, low-intensity stimulation: r= 0.314, p= 0.930; ‘20/80’, high-intensity stimulation: r= 0.315, p= 0.918; ‘80/20’, low-intensity stimulation: r= 0.607, p= 0.018; ‘80/20’, high-intensity stimulation: r= 0.408, p= 0.360; ‘50/50’, low-intensity stimulation: r= 0.417, p= 0.324; ‘50/50’, high-intensity stimulation: r= 0.417, p= 0.324).

## Discussion

The present study investigated neural processes underlying perceptual decisionmaking in the context of pain. We found that activity in the DLPFC reflected a direction-specific decision-making bias (as indexed by a change in starting point). While a bias towards low-intensity pain was positively correlated with DLPFC activity, a bias towards high-intensity pain showed a negative correlation with activity in the DLPFC (i.e. expectations of high-intensity pain induced less bias towards high pain judgments the more the DLPFC was engaged prior to stimulus application). This anticipatory DLPFC activity during the expectation of a high-intensity stimulation was linked to increased activation in the PAG during stimulus receipt. Changes in sensory processing (as indexed by a change in drift rate) were related to a heightened signal level in the amygdala and an increase in its functional connectivity with the PAG.

Using a similar modeling approach to the one we adopted here but employing affectively neutral stimuli, previous studies had linked a shift in starting point to increased activity in the DLPFC (Mulder et al., 2012; Nagano-Saito et al., 2012). Our data obtained using affectively laden stimuli confirm these previous observations but paint a more nuanced picture. In line with these previous findings, anticipatory DLPFC activity in our study scaled positively with the shift in starting point towards low pain when participants expected a low-intensity stimulation (Fig. 2). However, a *negative* relationship was found when high-intensity pain was expected (Fig. 2), albeit in a slightly different part of the DLPFC. The combination of these two findings challenges the notion of a direction-insensitive involvement of the DLPFC in bias implementation, and also suggests that its involvement depends on the nature of the expected stimulus. The DLPFC is known to be pivotal for the down-regulation of pain (Seminowicz and Moayedi, 2017) and more specifically for initiating top-down modulation prior to stimulus encounter (Krummenacher et al., 2010). Our findings are compatible with such a role in preparing the organism for subsequent stimulus encounter which is further supported by an investigation into the temporal profile of DLPFC activity in perceptual decision-making showing that activity peeked during stimulus anticipation not during stimulus receipt (Cardoso-Leite et al., 2014). Together, our results therefore provide further evidence for a role of anticipatory DLPFC activity in buffering against a bias towards high pain (or ‘keeping pain out of mind’; Lorenz et al., 2003) and supporting a bias towards low pain in healthy individuals.

Given our *a priori* hypotheses, our investigation strictly focused on the role of the DLPFC in (biased) decision-making and its link to model parameters. This is not meant to ignore the fact that the DLPFC - which spans a large part of the prefrontal cortex - is also pivotal to a number of other cognitive functions, as also evident from our analyses of the group main effects of the anticipation and stimulation period (see Supplementary Tables 1 and 2). Note, however, that our regression analyses were set up to ensure that our reported effects related to the model parameters are orthogonal to the group main effects (the regressor representing the individual participants’ model parameter only explains variance not captured by the main effect).

Research into cognitive-affective pain modulation has established that the DLPFC is embedded into a wider network of brain regions which complement its modulatory influence (see Atlas and Wager, 2014 for review). In line with this notion, we found increased activation in the PAG during stimulus *delivery* when DLPFC engagement had been high during stimulus *anticipation* (Fig. 3). The cluster was predominantly located in the ventrolateral part of the PAG (vlPAG) which has been implicated in endogenous analgesia (Bandler and Shipley, 1994 for review; Koutsikou et al., 2015; Ozawa et al., 2016; Tovote et al., 2016) and is connected with affective and evaluative prefrontal regions including the DLPFC (An et al., 1998; Floyd et al., 2000; Stein et al., 2012; Faull and Pattinson, 2017). DLPFC and PAG are key nodes of the descending pain control system that implements opioid-mediated expectancy-related pain modulation and most prominently placebo analgesia (Atlas and Wager, 2014; Krummenacher et al., 2010; Stein et al., 2012). Engagement of this system is commonly described as eliciting a direct influence on dorsal horn nociceptive processing. However, our results additionally suggest a link to biased decisionmaking (change in starting point) - a process that is believed to be different from sensory processing (change in drift rate). While a shift in starting-point is implemented at the time-point when relevant information becomes available (i.e., prior to stimulus delivery), the drift rate is tied to the processing of the stimulus itself. Because the drift-diffusion model used here does not allow for single-trial estimates, our data is not suited to probe the functional relevance of the observed DLPFC and PAG involvement in more detail. One could, however, speculate that prior information leads to a shift in starting point at the time the information becomes available (as reflected in DLPFC engagement) - but it is implemented through altered sensory processing (e.g., as implemented via antinociceptive processing reflected by involvement of the PAG) when the stimulus is applied. To explore this possible explanation, we tested whether the shift in starting point was correlated with altered drift rate in either of the conditions. Although these analyses only revealed a significant result for one of the six conditions (namely, the delivery of low-intensity stimuli when high-intensity stimuli had been expected), all comparisons showed positive correlations with medium to high effect sizes. This highlights that although model comparison in our previous study had established a shift in starting point as the dominant bias (Wiech et al., 2014), both types of bias are not mutually exclusive and further research is needed to explore whether they can operate in concert to bias perception.

Of note, significantly increased PAG activity during stimulus processing was only found when the expected high-intensity stimulation was applied but not when the high-intensity cue (‘80/20’ cue) was followed by a low-intensity stimulation or when participants had been cued towards low pain. In principle, PAG engagement could depend on expected outcome (e.g., it mainly occurs when the expected stimulation intensity is applied, irrespective of the intensity of stimulation), on intensity of stimulation (e.g., it mainly occurs when a high-intensity stimulation is applied, irrespective of the stimulation intensity expected) – or a combination of the two factors (e.g., it mainly occurs when the stimulation is expected and of a high intensity). Our findings seem to support the latter and suggest that PAG involvement during stimulus delivery might be most prominent in scenarios in which antinociceptive counter-regulation – one of the key functions of the PAG (Heinricher and Fields, 2013) – is required due to high intensity stimulation and facilitated through correct prior knowledge.

A second key finding of our study is the link between an increase in drift rate when high-intensity stimulation was unexpectedly received, the engagement of the amygdala (Fig. 4) and an increase in its functional connectivity with the PAG (Fig. 5). The unexpected delivery of a high-intensity stimulus can be regarded as the ‘worst case scenario’ in our paradigm and should evoke an aversive prediction error (PE). A multitude of studies have implicated the amygdala in aversive PE processing, showing that amygdala neurons respond preferentially to unexpected aversive stimuli (Belova et al., 2007; Johansen et al., 2010; Klavir et al., 2013; McHugh et al., 2014; Ozawa et al., 2016). Our observation of heightened amygdala activation to unexpected delivery of high-intensity noxious stimuli is thus supported by a large body of animal data and extends previous findings by demonstrating a link between PE-related amygdala activation and a change in drift rate. An increase in drift rate is often interpreted as accelerated sensory processing during evidence accumulation. From an evolutionary perspective, ‘fast tracking’ of incoming information seems adaptive when strong aversive input occurs unexpectedly, as only immediate changes in behaviour (e.g., escape or attack) may prevent further harm. Based on findings which showed that impaired amygdala functioning abolishes the heightened response to fear-related stimuli in primary sensory brain regions (Hadj-Bouziane et al., 2012; Rotshtein et al., 2009; Vuilleumier et al., 2004), the amygdala has been proposed to prioritize processing of emotionally relevant stimuli through gain control in these areas (Chen et al., 2014; Pourtois et al., 2013). Our data seem not to support this notion. Although activation in the left amygdala showed a positive correlation with change in drift rate during unexpected high-intensity stimulation (Fig. 4), we neither found evidence for altered processing in brain regions implicated in nociceptive processing (including thalamus, posterior insula, primary or secondary somatosensory cortex) nor a change in functional connectivity between the amygdala and these brain regions.

Recent investigations into the neural network underlying fear learning and its influence on sensory processing have focused on amygdala interactions with another structure – the PAG. Unexpected aversive stimuli generate a PE response in the PAG that serves as a teaching signal to drive learning and fear-related plasticity (Johansen et al., 2010; McNally et al., 2011). Notably, PE signaling in the PAG has recently also been demonstrated in the context of pain in humans (Roy et al., 2014). Here, we found an increase in the crosstalk between amygdala and PAG that scaled with the increase in drift rate when high-intensity stimulation was unexpectedly applied (Fig. 5). The location of this response was clearly distinct from the above-reported DLPFC-related PAG response: while the latter was observed in the vlPAG, the amygdala-related response was located much more dorsally. Such a dissociation is interesting with regard to animal data showing that vlPAG is mostly involved in setting learning asymptotes and implementing behavioral adjustments such as freezing or conditioned analgesia (Koutsikou et al., 2015; McNally et al., 2011; Ozawa et al., 2016; Tovote et al., 2016) whereas dorsal PAG projections back to the amygdala seem to be pivotal in guiding amygdala sensitivity to unconditioned stimuli (Kim et al., 2013; Ozawa et al., 2016).

The lack of modulation in sensory brain regions and the involvement of the amygdala and PAG which are both key regions of affective processing cast doubt on the exclusive interpretation of an increase in drift rate as amplified processing in sensory brain regions. As previously pointed out by Mulder and colleagues (2014), brain activity related to differences in drift rate could also reflect collinear cognitive processes such as attention, motivation or preparation of motor responses. The involvement of specific brain regions would thereby depend on the type of information that is accumulated. An involvement of the regions such as the amygdala is in line with growing evidence showing that expectancy manipulations of pain are not necessarily reflected in brain regions involved in sensory processing (Zunhammer et al., 2018) but might be reflected in regions associated with affective processing (Atlas and Wager, 2014). An increase in drift rate might therefore reflect the fast propagation of information within a system that ensures swift responses to impending threat, potentially including counter-regulatory processes. With amygdala responses to threat in less than 100 msec (Méndez-Bértolo et al., 2016) which arise prior to conscious perception (Bastuji et al., 2016) and connections with key regions of behavioural responses to threat, the amygdala is ideally suited to serve this function.

## Conclusions

Taken together, our results emphasize the relevance of affect-related considerations and accompanying cortico-brainstem interactions when investigating the neural basis of perceptual decision-making. Additional studies are needed to further explore the role of stimulus valence including its link to motivational aspects and learning which embed perceptual decisions into the context of the individual’s priorities. Such integration promises a novel and more comprehensive view on perceptual decision-making.

## Acknowledgements

K.W. was supported by an MRC UK New Investigator grant (MR/L011719/1). JZ is currently funded by the Research Foundation Flanders (FWO) and has been supported by the ‘‘Asthenes’’ long-term structural funding (METH/15/011) - Methusalem grant by the Flemish Government. J.W.S. Vlaeyen was supported by the Odysseus grant “The Psychology of Pain and Disability Research Program” funded by the Research Foundation Flanders, Belgium (FWO Vlaanderen, Belgium), and is currently supported by the “Asthenes” long-term structural funding Methusalem grant by the Flemish Government, Belgium (METH/15/011). F.T. was supported by KU Leuven Research Council Grant GOA/15/003 and the Fund for Scientific Research-Flanders (Grants G.0534.09N and G.0806.13). J.V. was supported by the Belgian National Science Foundation grant #1658303. I.T. wishes to acknowledge support from the MRC (UK) and Wellcome Trust. The Wellcome Centre for Integrative Neuroimaging is supported by core funding from the Wellcome Trust (203139/Z/16/Z).

